# multiScaleAC: Cell-Cell interaction with Moran’s I as a function of kernel bandwidth

**DOI:** 10.64898/2026.07.14.738306

**Authors:** Alex C Soupir, Mitchell T Hayes, Brandon J Manley, Xuefeng Wang, Lauren C Peres, Julia Wrobel, Brooke L Fridley

## Abstract

Over the last decade, spatial transcriptomic technology has transformed our understanding of tissue architecture including cell-cell interactions within the tumor immune microenvironment. A specific use-case of increasing interest is leveraging the spatial statistical relationship of genes whose protein products are known to be involved in ligand-receptor interactions. One methodological limitation of this approach has been the requirement to choose one radius around a cell as a parameter that can come with selection biases. Rather, interactions between cells vary in strength across a range of spatial scales that single-radius choice may miss. To fill this gap we developed ‘multiScaleAC’ to extended Moran’s I, a correlation measure that accounts for locations of values, by employing a Gaussian kernel applied to locations and varying the bandwidth parameter *h*. The resulting Moran’s *I*(*h*) then can be compared between samples using functional data analysis. In the current study, we used simulations to show that our framework has well controlled Type I error due to the use of permutations for assessing significant interactions. We also demonstrate that ‘multiScaleAC’ has high statistical power to identify a significant interaction when a true interaction is simulated (1.00 at bandwidths greater than 5) and increasing power as bandwidth increases when negative interaction is simulated. We found ‘multiScaleAC’ largely captures similar significant ligand-receptor profiles in 8 Visium samples of colon tissue using the same bandwidth as ‘spatialDM’ but without removing low-weight spots from the weight matrix (73.7% - 87.2%). Applying the ‘multiScaleAC’ framework to our previous single-cell spatial transcriptomics data (*COL4A1*-*ITGAV* in the stromal compartment of clear cell renal cell carcinoma) followed by functional principal component analysis, we found functional principal component 1 to represent global interaction elevation/depression. Associating functional principal component 1 scores with immunotherapy exposure showed significantly higher scores in stromal tissues exposed to immunotherapy than those naïve to immunotherapy, indicating an overall higher interaction of cell expressing *COL4A1*-*ITGAV*. These findings recapitulate our previous study while reducing bias in neighbor selections. We believe this is the first study to apply a functional extension of Moran’s I in combination with functional data analysis to understand cell-cell interaction over spatial scales.

## Introduction

The number of molecular studies involving spatial technologies like spatial protein imaging and spatial transcriptomics are drastically increasing. A literature search of PubMed reflects the accelerated use of these platforms, with 2,598 publications in 2023, 3,573 in 2024, and 5,657 in 2025 (pubmed.ncbi.nlm.nih.gov - accessed 26 May 2026) [1]. Along with the high volume of studies, the spatial resolution of technologies has rapidly advanced from region of interest (e.g., laser capture microdissection, Nanostring GeoMx), to single cell resolutions with multiplex immunofluorescence imaging, to subcellular localization of transcripts (e.g., Nanostring CosMx SMI, 10X Xenium) [2–6]. As resolution increases, so does the ability to use the data to form better understandings of the underlying biology. An insight that is provided greater power by this increase in spatial resolution is being able to assess cell-cell interactions using the location in which the molecules were measured at [7]. Highly multiplexed protein and spatial transcriptomics that are fluorescence-based allow for many ligand-receptor pairs or complexes to be profiled in the same tissue while also capturing the spatial locations which is a major advantage over traditional single-cell or laser capture microscopy studies. As a result, methods beyond simple correlation are needed to appropriately handle the added dimensionality of the data.

Existing computational approaches designed for identifying global cell-cell interactions have limitations. Spatial autocorrelation measures from ecology such as Moran’s I handle the spatial component by spatially lagging values before correlation [8–10]. However, current frameworks evaluate this relationship at only a single value, utilizing either a fixed Gaussian bandwidth or a fixed radius to define cellular neighborhoods [8, 11]. Selecting a single scale introduces subjectivity and bias into the analysis either through the selection process (i.e., selecting the radius that provides greatest differences between groups) or the information that is not captured (i.e., a single radius or bandwidth will leave out information at the other radii or bandwidths). Recently, this has been addressed in single-cell spatial clustering analyses with our package ‘mxfdà which computes spatial summary functions at a range of radii and uses the full curve (i.e., Ripley’s K by radii) as a feature for each sample but has not been addressed for ecology-based cell-cell interaction measures [12].

To overcome these limitations in cell-cell interactions and maximize sensitivity, we introduce ‘multiScaleAC’, an expansion of bivariate Moran’s / framework that models spatial autocorrelation as a continuous function of distances. By using a range of bandwidth parameters (*h*), we convert the static Moran’s *I* measure to a continuous interaction curve *I*(*h*). This interaction curve captures more information about the microenvironment than a single bandwidth or radius by including bivariate Moran’s I at many bandwidths [13]. We demonstrate that this multi-scale autocorrelation value has high power to robustly pick up significant interactions across varying cellular cluster sizes and spatial configurations while the false positive rate is fully controlled due to permutations. These *I*(*h*) curves can then be analyzed using functional data analysis (FDA) methods, such as functional principal component analysis (FPCA), to identify distinct spatial phenotypes across clinical cohorts.

## Methods

### Bivariate Moran’s *I* for estimating ligand-receptor interactions

Bivariate Moran’s *I* is an estimator of spatial autocorrelation. For two genes, it aims to assess the relationship between the expression of gene 1 at a location and gene 2 at the surrounding locations (spatially lagged). The estimate is formalized as:

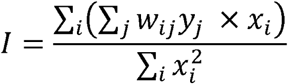

where *w_ij_* is a spatially-informed weight between cells *i* and *j*, and *x* and *y* are genes 1 and 2, respectively.

In Moran’s initial description of the function weights were binary [9], with weight of 1 for points that were considered spatial neighbors and 0 for those that were not. This results in unbound *I* values. However, with the application of row-standardization of weights (summing to 1), the values of *I* are then bound by -1 and 1 [14]. When values trend together spatially, *I* approaches 1 and when values trend opposite spatially the measure of *I* approaches -1. Further, row-standardization of the binary weights make all *I* cells contribute the same weight towards the final *I*. For point patterns representing biological contexts, points that are closer should contribute more to interactions between cells when a binary weight is no longer sufficient.

### Gaussian kernel as cell-cell weight matrix

The weights between cells can, in theory, be anything that describes their spatial relationship. Previously, researchers used a Gaussian kernel applied to point/cell locations as weights after removing low kernel density estimates (KDEs) through threshold filtering [11]. This produces a single estimate of accounting for distance. The use of a Gaussian kernel applied to cell locations for their spatial weights provides greater weight to immediate neighbors than those which are further away. Conversely, the use of fixed neighbors with KNN would not take distance into account resulting in cells in a sparse sample contributing as much to the ‘interaction’ measure as those in dense sample.

### Functional extension of bivariate Moran’s I

Using a Gaussian kernel applied to the point/cell locations still relies on the selection of a bandwidth that is ‘right’ for each data. We extend this Gaussian kernel framework to capture *I* from a range of bandwidths to generate a functional curve that represents multis-scale spatial autocorrelation, or ligand-receptor interaction, which we call multiScaleAC. The weight matrices are then represented by *w*(*h*) rather than w:

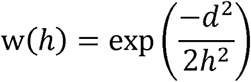

where *d* is then *n* × *n* distance matrix for a sample with *n* cells. By using this definition of weights, the bivariate Moran’s *I* becomes I(*h*) that represents the interaction between gene 1 and gene 2 at varying bandwidths, or spatial scales:

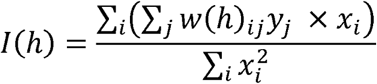

This *I*(*h*) functional curve removes the need to select a radius or bandwidth parameter that introduces bias in analyses. To generate a null distribution of *I*(*h*) under complete spatial randomness, values of are subtracted from the observed *I*(*h*) to create a measure of spatial autocorrelation are permuted. The average of the permutations (analytically 0 for no spatial association) that removes the dependency of cell locations. Further, the distribution of curves provides an empirical value for significance. Bandwidth-wise significance is calculated by comparing the observed value to the permutation distribution at each bandwidth, counting the number of permutations greater than the observed (one tailed) and dividing by the total number of permutations (p value). We also calculate the significance of the full curve using supremum which finds the bandwidth with greatest deviation from the mean curve for each permutation and compares it to the largest deviation of the observed.

### Datasets

We used two platforms to demonstrate ‘multiScaleAC’’s cell-cell interaction framework: 10x Visium and Nanostring CosMx 1k. The 10x Visium samples were intestinal from the ‘SpatialDM’ Python module (raw data available: GSE158328) while the CosMx 1k samples were from our previous work of clear cell renal cell carcinoma (ccRCC; raw data available on Zenodo) [7, 11, 15, 16]. Clear cell renal cell carcinoma samples were from matched tumor and stroma core (adjacent tissue) on 3 tissue microarrays (TMAs). Intestinal samples (n=8) were from an adult colon (A1 and A2), fetal whole colon 12 and 19 weeks post conception (A3 and A4, respectively), and whole small intestine (A6 and A7) and whole colon (A8 and A9) 12 weeks post conception.

’CellChat’ [17] database was exported from the ‘SpatialDM’ python module and ‘CellTalk’ [18] database were downloaded from GitHub (https://github.com/ZJUFanLab/CellTalkDB). When there were multiple genes that formed either a ligand or receptor complex, the expression was averaged. If the data set did not contain both a gene from the ligand complex and a gene from the receptor complex, the ligand-receptor was discarded from analyses. Further, due to the Visium and CosMx data being sparse, if a ligand-pair didn’t show expression in 3 or more cells/spots [11], that ligand-receptor pair was dropped.

The ‘SpatialDM’ global *I* and ‘multiScaleAC’ were applied to the Visium intestine data with the ‘CellChat’ gene lists. For ‘SpatialDM’’s implementation, we attempted to replicate the parameters used in the published study with a bandwidth of 75 and a kernel density estimate (KDE) filtering threshold of 0.2 (following their ‘differential_test_intestine.ipynb’ notebook). Significance of global bivariate Moran’s I was calculated with 1000 permutations and an FDR threshold of 0.05. To compare to our approach, we used Gaussian kernel values as weights without applying the low filtering threshold (bandwidth of 75).

### Multi-scale functional analysis of interaction curves

Because each ligand-receptor interaction with ‘multiScaleAC’ yields a full interaction curve over a range of bandwidths rather than a single interaction value, we used functional data analysis to compare interaction profiles between CosMx SMI sample groups: treatment naïve and immunotherapy (IO) exposed stromal tissues. Bandwidths from 0 to 500 were calculated, and bandwidths from 10 to 250 were used for functional data analysis. This reduction in range removes the unstable region close to 0 as well the region at higher bandwidths where values of I(h) do not change drastically. The ‘mxfdà package was used to perform functional principal component analysis on the curves [12]. Wilcoxon rank-sum test was performed on FPC1 to show discrimination between treatment naïve and treatment exposed stroma FOVs. Code for calculating Moran’s I functional curves can be found at https://github.com/ACSoupir/KernelCommunication.

### Simulations for Type 1 error rate and power

To quantitatively validate the statistical sensitivity and false positive rate of our functional bivariate Moran’s *I*(*h*) computational framework, we constructed *in silico* spatial interaction scenarios. Synthetic spatial environments were generated using the ‘scSpatialSIM’ R package [19], which allowed us to construct controlled interaction structures while mirroring the authentic expression distribution of actual biological tissues, specifically, gene expression for ligands/receptors and cell/point densities. We explicitly reconstructed two primary spatial architectures: a Visium environment with gridded spots (from SpatialDM sample A1), and a high-resolution CosMx SMI architecture with random placement (RCC3-1). The simulated point pattern intensities, ligand/receptor expression, and 0-expressing dropouts were derived from the real samples can be found in **Table 1**.

**Table 1:** Empirical expression profiles used in simulations to assess false positive (Type 1) error. No spatial structure was simulated.

To quantify the power of our method to pick up true spatial interactions between ligand and receptor hotspots, we again used ‘scSpatialSIM’ [19] to simulate spots/cell densities based on A1 from Visium and RCC3-1 from CosMx SMI. Visium spots and CosMx SMI cells expressing the ligand or receptor were simulated at different cluster sizes and different shifts (**Table 2**). Meaning, regions with simulated expression of a receptor was either the same as the ligand or shifted to a Dirichlet vertex (full separation of expressing spots or cells). The size of hotspot expressing the ligand and receptor was either small or large.

**Table 2:** Simulation parameters for assessing power.

## Results

Comparison of ‘SpatialDM’ to our unbiased kernel approach in ‘multiScaleAC’ was performed at a bandwidth of 75 for both methods, with ‘SpatialDM’ additionally filtering kernel density estimates less than 0.2. The number of ligand-receptor pairs that met the minimum spot thresholds for genes in the data and minimum spots with expression ranged from 1169 (A1) to 1266 (A4; **Table 3**; **Figure 1**). Both methods have high overlap of the ligand-receptor pairs identified as having significant spatial relationships (73.7% - 87.2%). In 5 of the 8 10x Visium intestinal samples, ‘multiScaleAC’ identified more pairs significant while in 3 samples ‘SpatialDM’ identified more ligand-receptor pairs significant.

**Figure 1:**
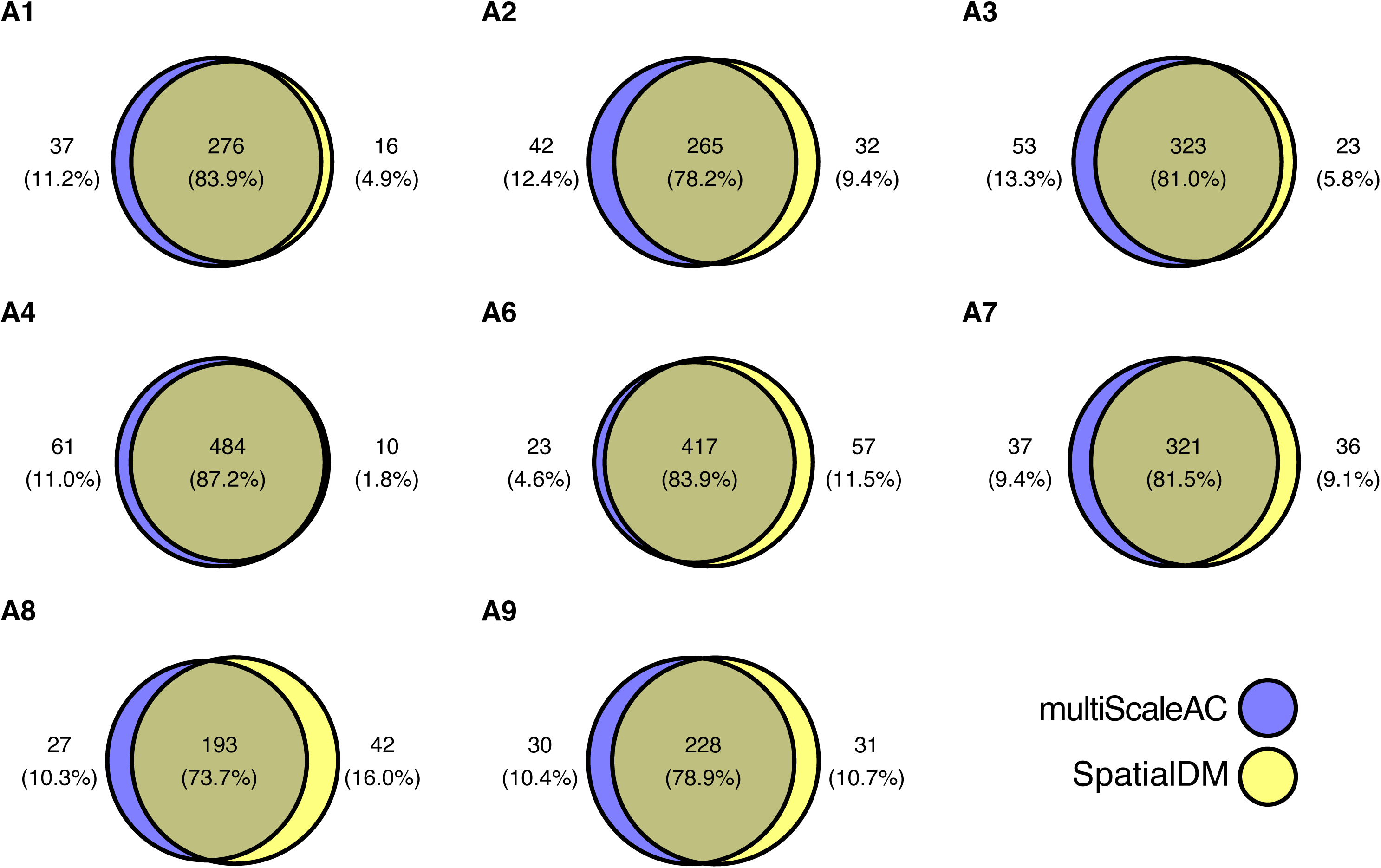
Intersection of ligand-receptor pairs identified as significant with ‘SpatialDM’ and ‘multiScaleAC’. Intestinal samples profiled with 10X Visium (GSE158328) had cell-cell interactions profiled for ligand and receptor pairs found in ‘CellChat’ database to identify the amount of overlap between the methods at a bandwidth of 75. ‘SpatialDM’ uses a global scaling for their standardization and minimum weight threshold of 0.2 while ‘multiScaleAC’ uses row-wise standardization with no minimum weight threshold.

**Table 3:** Number of ligand-receptor pairs that were identified as significant in SpatialDM, KernelCommunication, niether, or both methods in Visium samples from intestine with a bandwidth of 75. Both methods used permtuations to determine significance (FDR < 0.05).

To determine the Type I error rate of ‘multiScaleAC’, we simulated varying expression levels of ligand and receptor “genes”, and randomly placed the positive points/cells in coordinate systems based on real Visium and CosMx SMI samples (**Table 1**). All simulations were around 5% Type I error rate, ranging from 3.2% (Scenario 11 on CosMx *TFGB1*-(*TBFBR1*+*TGFBR2*) ligand-receptor expression) to 7% (Scenario 4 on Visium with the most 0s; **Table 1**). This is demonstrated across all bandwidths except for those close to 0 which fail to capture points that are near enough to provide weight (**Figure 2**). The Type I error also is stable from low to high bandwidths due to the use of permutations to generate the underlying null distribution for complete spatial randomness.

**Figure 2:**
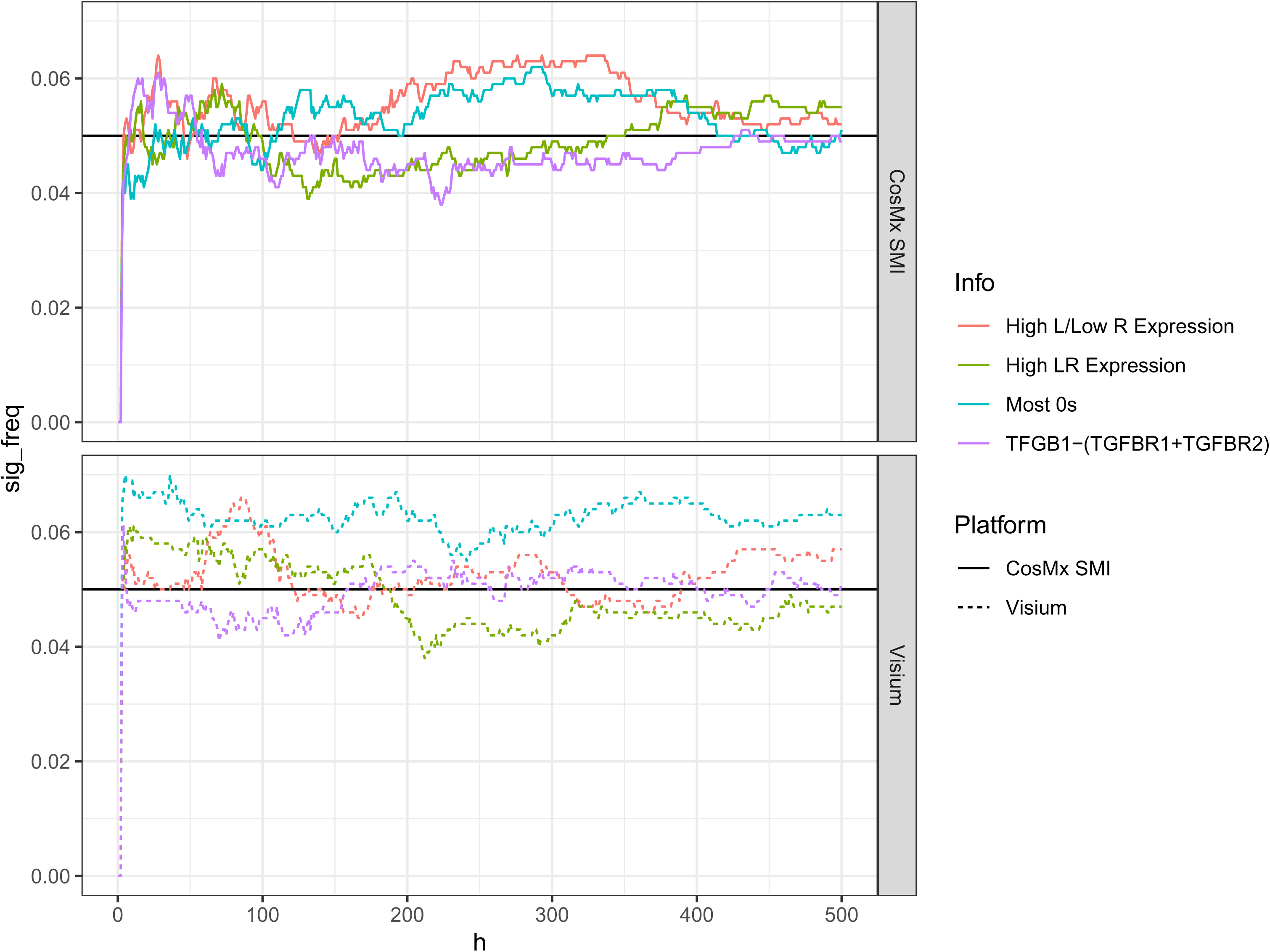
Type I error rate of ‘multiScaleAC’ for simulations based on real Visium intestine and CosMx 1K clear cell renal cell carcinoma data.

The ability to pick up cell-cell interaction when a strong positive signal is simulated is critical for detecting real ligand-receptor interactions. We simulated gene expression hotspots with shifts between ligand and receptor hotspots. **Figure 3** demonstrates one such scenario where gene 1 (ligand) and gene 2 (receptor) were shifted to Dirichlet vertices (all 8 scenarios described in **Table 2**). Under scenarios where the location is the same for the ligand and receptor (true spatial relationship; no shift), our ‘multiScaleAC’ identifies significant interaction, where the observed Moran’s I value is more extreme than the permuted null distribution, at all but low bandwidths (other points/cells are not contributing to the weights; **Table 4**; **Figure 4**). At all bandwidths greater than 5, 100% of simulations had significant interaction detected regardless of small or large clusters when true spatial relationship was simulated.

**Figure 3:**
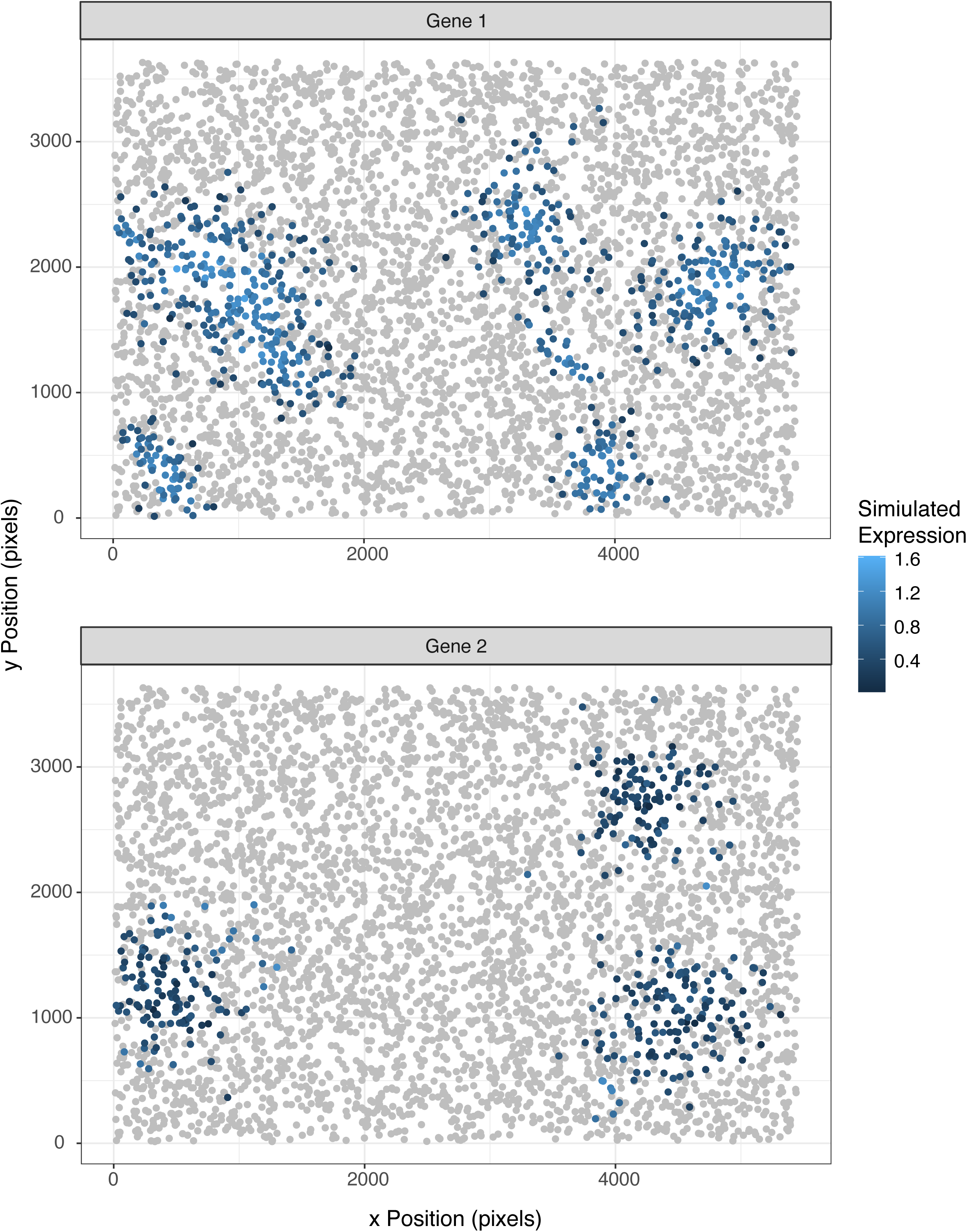
Example CosMx SMI simulation demonstrating the shift of gene 2 hot spots from gene 1 hotspots. This represents full segregation of ligand-receptor interactions.

**Figure 4:**
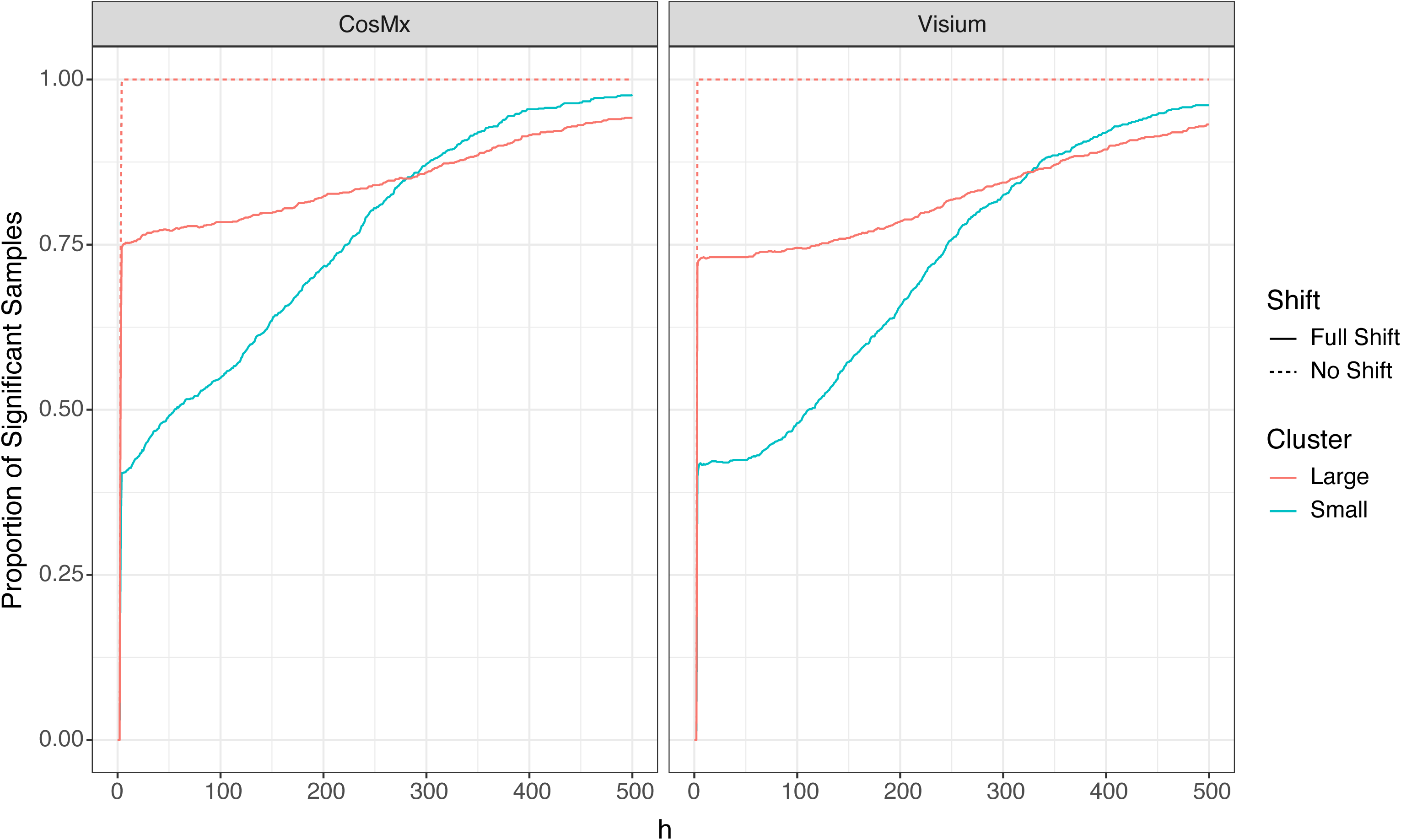
Power of our method to pick up significant ligand-receptor interaction.

**Table 4:** Power of Moran’s I in varying simulation scenarios using different kernel bandwidths.

When a full shift to Dirichlet vertices was done for the receptor expression centers, ‘multiScaleAC’ identified less interaction than true spatial relationship, yet respectable (∼40% at the same bandwidth of 5; **Table 4**; **Figure 4**). Large clusters under full shift of the receptor location showed higher power at lower bandwidths than the smaller clusters for both technologies. Both large and small cluster sizes of the gene expression show increasing power as bandwidth increases due to the bandwidth providing reach to the neighboring receptor hotspots.

In our previous work, we identified significantly increased spatial autocorrelation of *COL4A1* and *ITGAV* in the stroma from patients who had received IO when compared to stroma from patients who have not yet been given IO. Applying our multiScaleAC interaction from *h* = 5 to *h* =250 for each sample (**Figure 5**). The curves generated for framework to calculate the interaction between *COL4A1* and *ITGAV*, we extracted the those samples that were IO exposed follow a consistent trend of higher interaction at lower bandwidths and slow leveling off as bandwidth increases. There are less consistent, common interaction trends in the stroma from tumors that have not been exposed to IO (**Figure 5**).

**Figure 5:**
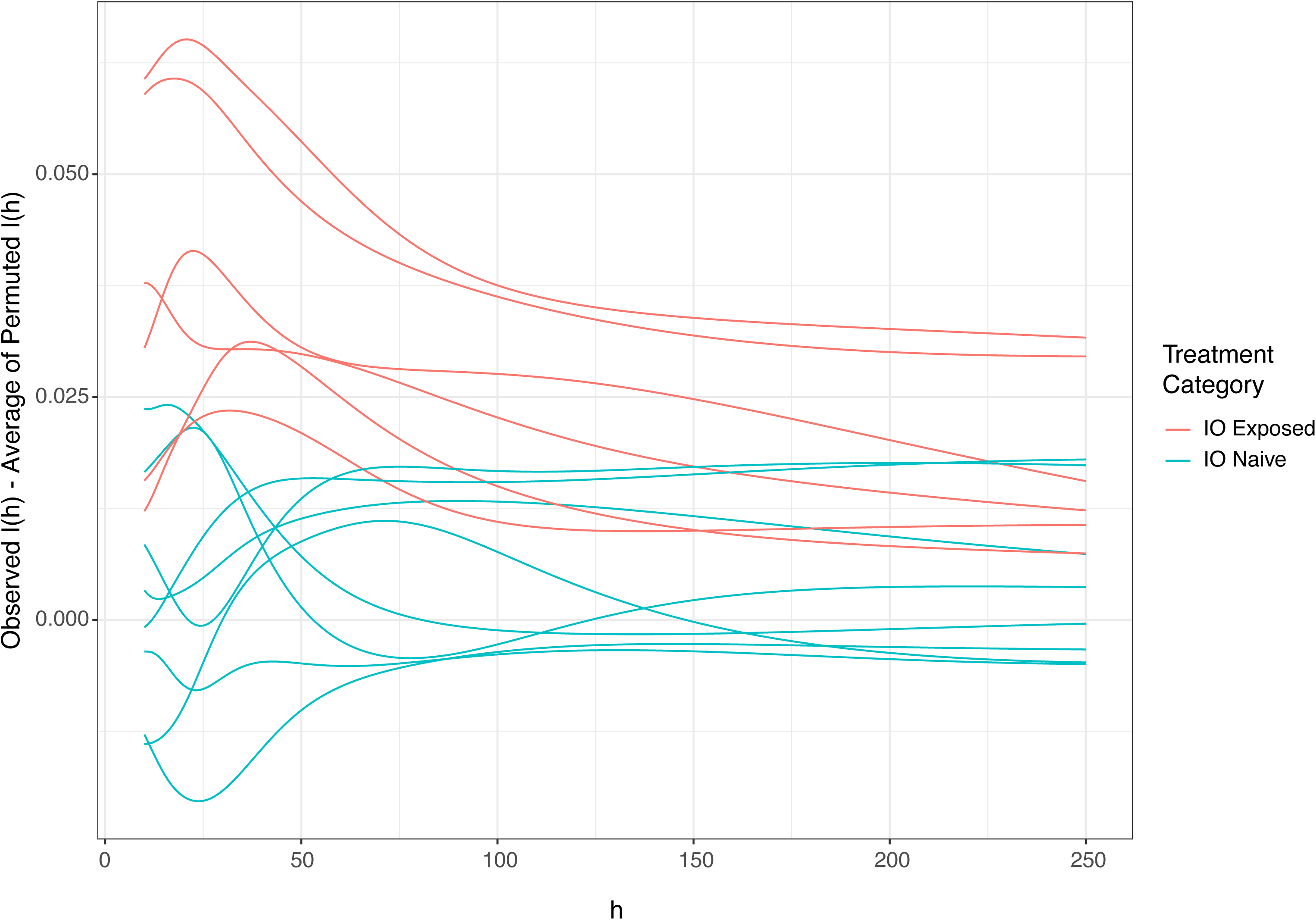
Interaction curves from ‘multiScaleAC’ for *COL4A1*-*ITGAV* across a range of bandwidths in immunotherapy exposed and immunotherapy naive stromal fields of view.

Applying functional data analysis methods like functional principal component analysis (FPCA) to the curves allows us to identify deviations from the average and assess differences between immunotherapy exposures. All stromal samples (n=?) from primary ccRCC produced interaction curves, so none were dropped. Three functional principal components were needed to represent 99.3% of variance (**Table 5**). Inspecting FPC1 (91.2% variance explained) shows direction of greatest variance is largely global across all bandwidths, with a positive FPC1 score being associated with elevated interaction (**Figure 6A**). FPC2 (6.5% variance explained) describes variation in the shape of the interaction curves rather than a uniform upward or downward shift. Specifically, it contrasts interaction at smaller and larger bandwidths: positive scores correspond to curves that are relatively lower than the mean at bandwidths below approximately 50 and relatively higher than the mean at bandwidths above 50, with negative scores producing the reverse pattern.

**Figure 6:**
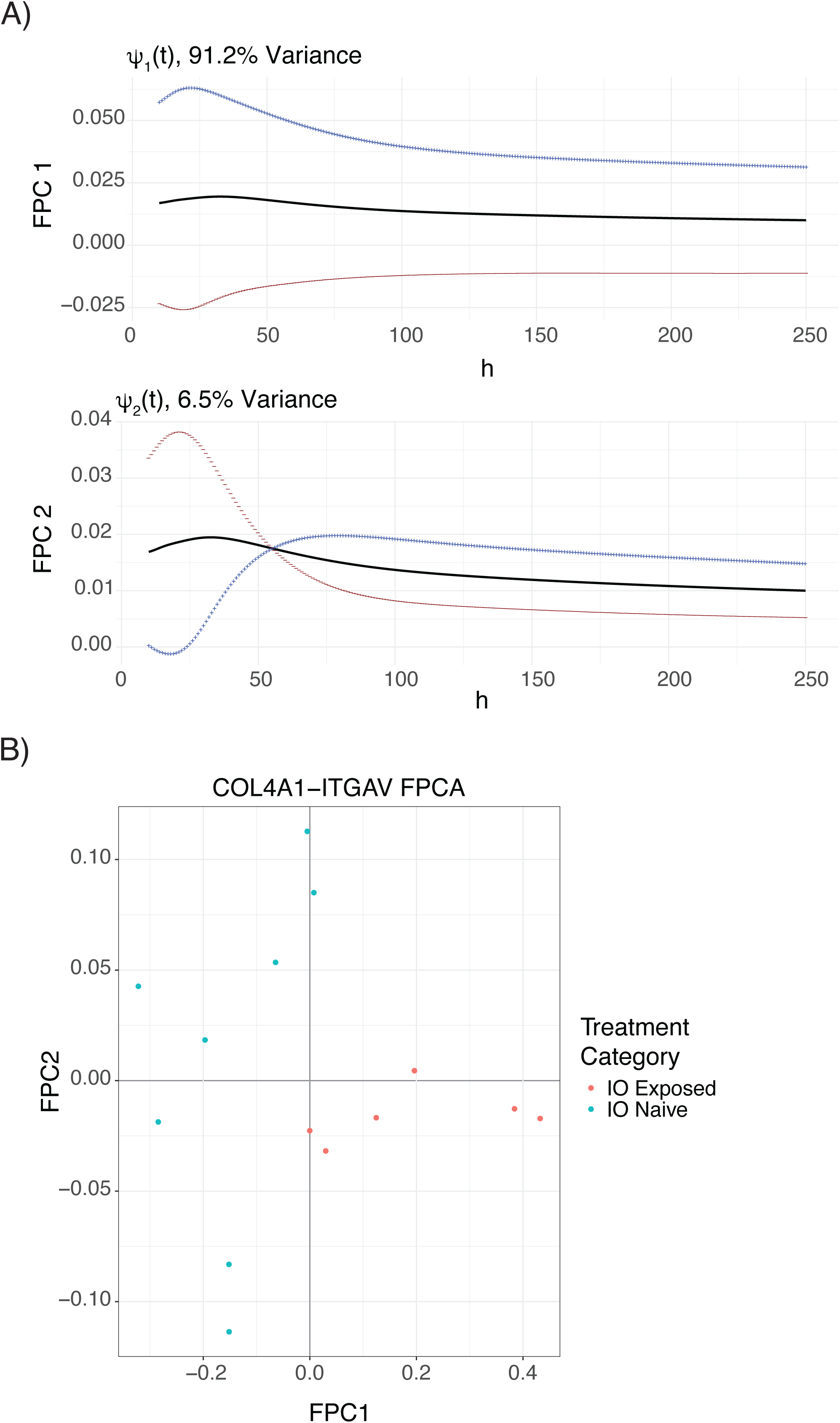
Functional principal component analysis (FPCA) of *COL4A1*-*ITGAV* interaction. A) shows the shape and 2 standard deviations (red and blue) from the mean (black) for what respective score represent. B) demonstrates separation of samples based on FPC1.

**Table 5:** Functional principal component scores. The first three principal component scores that represent 99% of the variance between samples.

Plotting FPC1 scores vs FPC2 scores shows how immunotherapy exposure clusters samples. The first FPC is significantly associated with immunotherapy exposure (Wilcoxon p = 0.0013; **Figure 6B**). The elevated scores associated with greater interaction between *COL4A1* and *ITGAV* are driven by samples exposed to immunotherapy (-0.002 to 0.432) while samples naïve to immunotherapy are lower (-0.322 to 0.008), related to less interaction than the average (**Table 5**). FPC2 variation is due to samples that were not exposed to immunotherapy (-0.11 to 0.11) while immunotherapy exposed samples have a much lower range (-0.03 to 0.004).

## Discussion

We set out to develop a multiscale Moran’s I spatial autocorrelation measure, Moran’s *I*(*h*), to minimize the bias of radius, bandwidth, or k-NN selection, and implement it into the R package ‘multiScaleAC’. We show that by using permutations to determine significance provides control of Type I error over different simulated expression and technology scenarios. We also show that when there is true simulated interaction, ‘multiScaleAC’ consistently demonstrates high statistical power to identify significant interaction, and when there are shifts in ligand-receptor expression hotspots the power of ‘multiScaleAC’ to identify significant interactions increases as bandwidth increases. Comparing ‘multiScaleAC’ using the bandwidth of 75 in colon Visium samples from ‘spatialDM’, we show high agreement of ligand-receptor interactions detected ranging from 73.7% - 87.2% agreement (**Figure 1**) [11]. The difference in the number identified as significant between ‘multiScaleAC’ and ‘spatialDM’ is likely a result of differences in standardization (’multiScaleAC’ uses row-standardization while ‘spatialDM’ uses global standardization) and ‘multiScaleAC’ not performing low-end thresholding for the Gaussian weights (’spatialDM’ removed weights less than 0.2) [11]. We also demonstrate that by applying ‘multiScaleAC’ to our previous single-cell spatial transcriptomic data (*COL4A1* and *ITGAV* expression in stromal fields of view) followed by functional principal component analysis (FPCA), the direction of largest variance (FPC1) is significantly associated with exposure to immunotherapy (p = 0.0013). These findings demonstrate that using a multi-scale spatial autocorrelation measure we can identify significant cell-cell interactions while minimizing bias.

Previous spatial studies focus on the use of a single value of radius, bandwidth, or near neighbor count which can be difficult to appropriately select without introducing bias [7, 11, 20–25]. Selecting a single radius can be based on prior biological knowledge but is challenging due to differences in protein diffusion in tissues. Further, k-NN means each cell/point has the same number of samples for the lagging weight of the receptor but loses spatial context which is influenced by the number of cells/points per unit area. By eliminating the need to select a single value, ‘multiScaleAC’ allows for more information from a range of spatial scales to be used in analyses. With the underlying structure of ‘multiScaleAC’ being modular, the distance measure and kernel can be swapped for something else that may better characterize the molecules. Further, the output can be easily incorporated with existing packages with ‘mxfdà’s ‘add_summary_function()’ making downstream functional data analysis methods easy to implement [12]. However, since the autocorrelation is being calculated at each bandwidth, it does take longer to compute.

In our previous work, we found that spatial autocorrelation between *COL4A1* and *ITGAV* was significantly higher in immunotherapy-exposed stromal tissues than in immunotherapy-naïve stromal tissues using a neighborhood defined by three nearest neighbors [7]. Here, the multiscale analysis extends that finding by showing that the increased spatial autocorrelation associated with immunotherapy exposure is not restricted to a single neighborhood definition but persists across a range of spatial scales. FPCA further separates two features of these interaction profiles: FPC1 captures differences in the overall magnitude of spatial autocorrelation across bandwidths, whereas FPC2 captures differences in curve shape by contrasting interaction at smaller and larger bandwidths. Thus, the multiscale approach provides evidence that the previously observed association is robust to the choice of spatial scale while also identifying variation in how the interaction changes as progressively more distant cells contribute to the measure.

While the current implementation of Moran’s *I*(*h*) in ‘multiScaleAC’ reduces subjectivity in radius, bandwidth, or k-NN selection and allows researchers to assess interaction profiles across spatial scales, it has its limitations. The main limitation of ‘multiScaleAC’is the use of permutations to remove the measure of complete spatial randomness and assess significance of the observed interaction. When calculating complete spatial randomness with permutations (random assigned locations of gene 2), the average should in theory be a value of 0 across all bandwidths regardless of the underlying spatial architecture. Though, to identify significant deviations from complete spatial randomness, the full distribution is currently required. Permutations in combination with the previously mentioned need for calculations of each bandwidth means that it can be very compute intensive. This provides future opportunities for code optimizations as well as possible identification of analytical mean and variance of the distributions, eliminating the requirement of permutations. Additional future work includes the inclusion and assessment of local Moran’s I(h), which will require multilevel FPCA to handle repeated measures, in addition to adding different kernels and distance measures to the ‘multiScaleAC’ package.

